# GluTrooper: a novel reporter mouse line for whole-brain imaging of glutamate dynamics

**DOI:** 10.1101/2025.11.11.686940

**Authors:** Gian Marco Calandra, Daniel P. Varga, Paul Feyen, Francesco Gubinelli, Benedikt Wefers, Peter Makra, Joshua James Shrouder, Andrea Cattaneo, Sumeyye Gökce, Wolfgang Wurst, Arthur Liesz, Jochen Herms, Lena F. Burbulla, Severin Filser, Nikolaus Plesnila

## Abstract

Glutamate is the primary excitatory neurotransmitter in the mammalian brain. However, tools to image glutamate dynamics in the whole brain with high spatial and temporal resolution are lacking. Therefore, we developed GluTrooper, a novel mouse line engineered for inducible and long-lasting expression of the genetically encoded glutamate sensor iGluSnFR3. GluTrooper mice crossed with Emx1-Cre lines demonstrated uniform and stable sensor expression in excitatory neurons of the cortex, hippocampus, and olfactory bulb. iGluSnFR3 expression remained stable for at least 12 months, enabling longitudinal observations of glutamate dynamics over extended periods. Using multimodal imaging in awake mice, we demonstrated the versatility of GluTrooper across multiple spatial scales: from mesoscale widefield cortical imaging to cellular resolution with two-photon microscopy. Moreover, during cortical spreading depolarization, bilateral whole-brain glutamate dynamics and contralateral cortical disinhibition were detected with high fidelity. Accordingly, the GluTrooper may open new avenues for the better understanding of glutamatergic neurotransmission in the mammalian brain.

## Introduction

Glutamate is the principal excitatory neurotransmitter in the brain, shaping the excitatory-inhibitory balance alongside GABAergic signaling [1, 2]. Hence, the ability to monitor extracellular glutamate *in vivo* is regarded to be a desirable tool for the better understanding of neural circuit function in both physiological and pathological conditions. Recent advances in genetically encoded fluorescent sensors, particularly the intensity-based glutamate sensing fluorescent reporter (iGluSnFR), have enabled direct, precise, high-resolution tracking of glutamate dynamics *in vitro* and *in vivo* [3–6]. While adeno-associated virus (AAV)-mediated delivery is the common technique for sensor expression in the brain, it presents several challenges for longitudinal imaging studies. AAV expression often develops slowly and can vary unpredictably over time, leading to patchy expression and potential off-target labeling, which complicates data interpretation. Additionally, strong continuous expression may result in aggregation and cellular toxicity [7, 8]. Moreover, AAV-mediated transfection requires a surgically invasive procedure, which may alter the cortical microenvironment. Fluorescent sensors delivered via AAV are also prone to photobleaching during repeated *in vivo* imaging sessions, further limiting signal stability and consistency. To address these limitations with the aim of achieving consistent and long-term glutamate imaging *in vivo*, we developed a ROSA26 knock-in mouse line expressing iGluSnFR3, referred to here as ‘GluTrooper’. Crossing this line with Emx1-Cre mice [9] resulted in broad expression of the sensor in the cell membrane of excitatory neurons throughout the cortex, hippocampus, and olfactory bulb. This reporter line enables stable, longitudinal *in vivo* monitoring of glutamatergic signaling across the brain in awake mice, under both physiological conditions and pathological events such as cortical spreading depolarizations (CSDs).

## Results

### Generation of a conditional knock-in mouse for targeted and longitudinal expression of iGluSnFR3

To enable the investigation of glutamatergic neurotransmission, we utilized a knock-in strategy targeting the well-characterized ROSA26 locus, which permits robust expression of transgenes while avoiding the interference of critical endogenous gene functions [10, 11]. The targeting construct included 5’ and 3’ homology arms (ROSA 5’HA and ROSA 3’HA) flanking the functional elements to ensure precise integration by homologous recombination (**Fig. 1A top**). The construct contained an AG slice promoter followed by an FRT-flanked sequence, a ubiquitous CAG promoter to drive high level expression, and a loxP-flanked puromycin resistance cassette (puroR) acting as a STOP signal in front of the iGluSnFR3 coding sequence. The presence of the STOP cassette prevents the expression of iGluSnFR3 until Cre-mediated recombination removes the intervening sequence, making this design vital for achieving conditional expression. Additional regulatory elements included the Woodchuck Hepatitis Virus Posttranscriptional Regulatory Element (WPRE) to enhance mRNA stability and translation efficiency, as well as human growth hormone (hGH) and SV40 polyadenylation signals to ensure proper mRNA processing and stability. The iGluSnFR3 protein is targeted to the plasma membrane via an N-terminal IgΚ secretion leader sequence and a C-terminal glycosylphosphatidylinositol (GPI) anchor, which promote efficient membrane localization [3]. The resulting mouse line (ROSA26-CAG-lsl-iGluSnFR3), named as GluTrooper, carries the iGluSnFR3 transgene in all cells but expresses it only upon Cre recombinase. Genotyping confirmed successful germline transmission and maintenance of the transgene through multiple generations, establishing a stable line for experimental use. To achieve pan-cortical expression, GluTrooper mice (**Fig. 1A top**) were crossed with Emx1-Cre mice [9] (**Fig. 1A bottom and B**). Given the cerebellum’s distinct developmental origin and lack of transcription factor Emx1 [12], it served as an internal control for regional iGluSnFR3 intensity quantification [13] (**Fig. 1C**). Regional analysis (**Fig. 1D**) revealed robust iGluSnFR3 intensity levels in the somatosensory cortex (**Fig. 1E**), hippocampus (**Fig. 1F**) and olfactory bulb (**Fig. 1G**) compared to cerebellar levels. Importantly, iGluSnFR3 expression showed no significant regional variation across developmental timepoints (**Fig. 1H-J**), making the GluTrooper a unique tool for mapping glutamatergic neurotransmission in both acute and longitudinal investigations.

**Figure 1.**
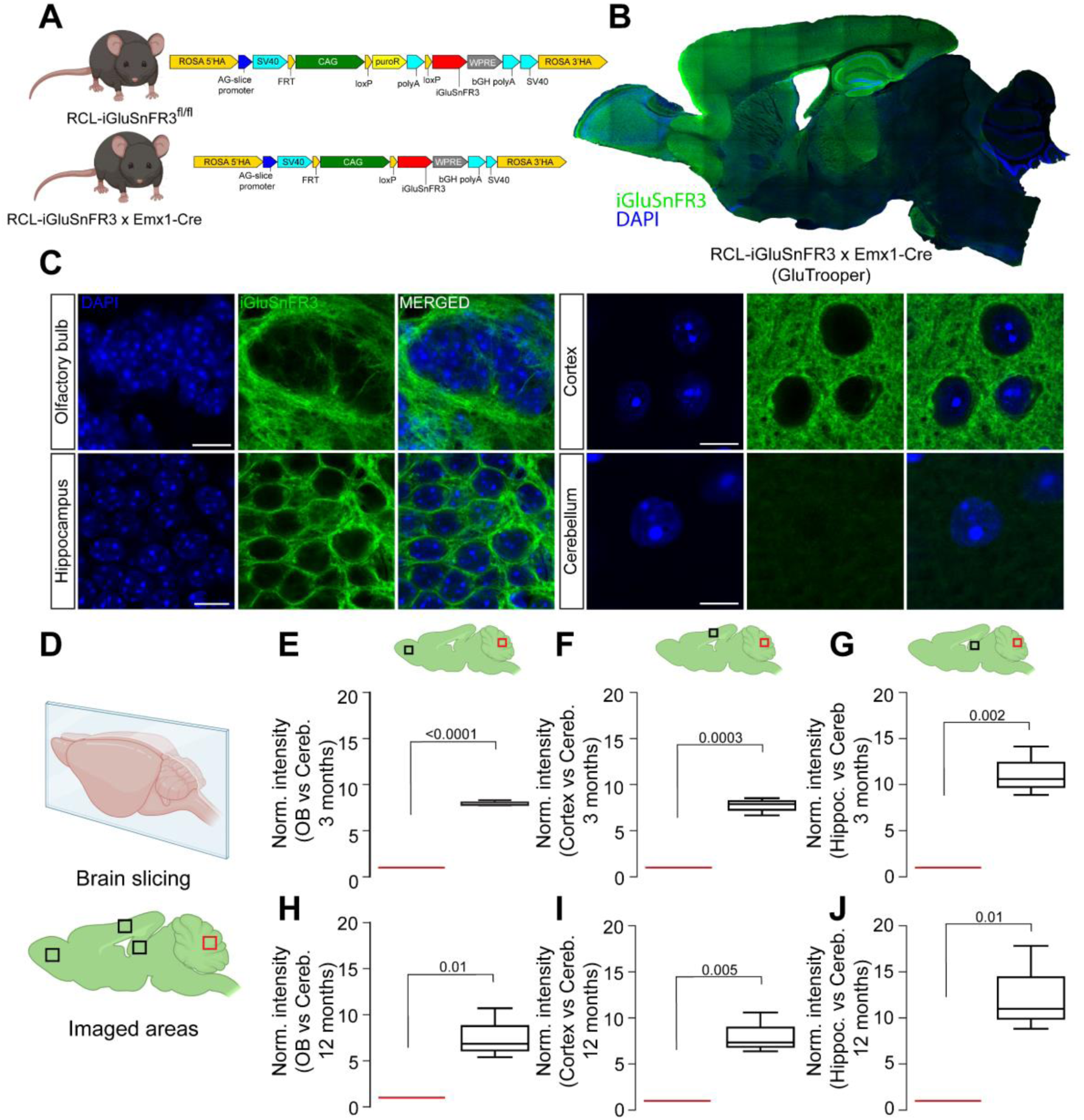
Generation of a conditional knock-in mouse for targeted and longitudinal expression of iGluSnFR3. **A)** Schematic illustrating the genetic design of the GluTrooper mouse line (top) and the resulting construct following crossbreeding with Emx1-Cre mice (bottom). **B)** Representative sagittal brain section showing widespread, pan-cortical expression of iGluSnFR3. **C)** Representative images from four distinct brain regions: olfactory bulb, hippocampus, cortex, and cerebellum (scale bars = olfactory bulb: 10 µm; somatosensory cortex: 10 µm; hippocampus: 10 µm; cerebellum: 5 µm.). Note the lack of iGluSnFR3 expression in the cerebellum due to lack of transcription factor Emx1. **D)** Schematic overview of the brain slicing strategy (top) and the four analyzed brain regions (bottom). **E-J)** Quantification of iGluSnFR3 fluorescence intensity in the olfactory bulb, the somatosensory cortex, and the hippocampus in comparison to the cerebellum at 3 and 12 months of age. Data are presented as normalized to cerebellar intensity levels (n = 3 mice per group, Student’s *t*-test). All data are displayed as median and the interquartile range (25th to 75th percentiles), whiskers reach to the minimum and maximum values of the distribution. Schematic illustrations are created with BioRender.com.

### Maintained glutamate specificity of the iGluSnFR3 sensor in GluTrooper mice

To assess iGluSnFR3 functionality in the GluTrooper line, we performed *ex vivo* glutamate imaging in acute brain slices (**Fig. 2A**). Fluorescent patches within the neuropil were clearly distinguishable and exhibited spontaneous fluctuations in iGluSnFR3 fluorescence (**Fig. 2B and Suppl. video 1**). To verify that the sensor iGluSnFR3 expressed in the GluTrooper line remains specific for the neurotransmitter L-glutamate, we applied the stereoisomers, D-glutamate and L-glutamate, at equimolar concentrations of 500 μM (**Fig. 2C**). GluTrooper brain slices displayed markedly different responses to these stereoisomers (**Fig. 2D**). Application of L-glutamate, the physiologically active stereoisomer, resulted in a substantial fluorescence increase of 91% (**Fig. 2E-F and Suppl. video 2-3,** L-glutamate vs D-glutamate, p=0.0008). In contrast, D-glutamate increased iGluSnFR3 fluorescence by only 3%, failing to elicit comparable changes in iGluSnFR3 fluorescence (**Fig. 2E and F**). These results demonstrate that the iGluSnFR3 sensor expressed in the GluTrooper line maintains identical glutamate specificity to the virally-delivered parental construct [3, 14, 15], enabling reliable visualization of spontaneous and evoked glutamate dynamics.

**Figure 2.**
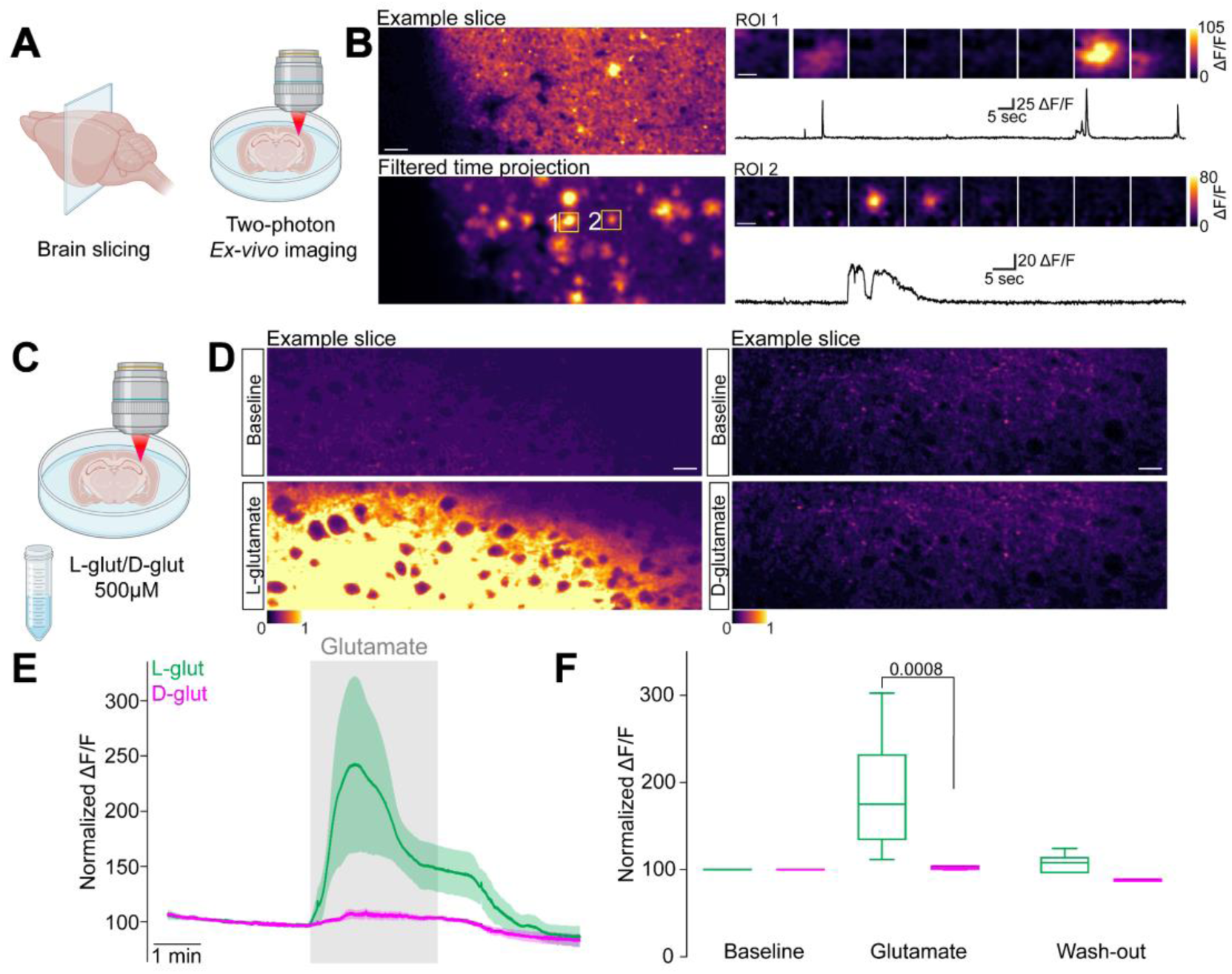
Maintained glutamate specificity of iGluSnFR3 sensor in GluTrooper mice. **A)** Schematic illustrating the *ex vivo* experimental approach. **B)** Raw time projection of an example slice (top) and corresponding filtered time projection (bottom); Scale bar = 20 µm. Note the various fluorescent patches identified within the neuropil. On the right, two example ROIs identified in the filtered time projection and corresponding activity; Scale bars: ROI1: 10 µm; ROI2: 5 µm. **C)** Schematic representing the experimental approach to test glutamate selectivity of the iGluSnFR3 expressed in the GluTrooper line **D)** Representative images showing detected glutamate fluorescent levels in presence of L-glutamate (left) and D-glutamate (right); Scale bars: 20 µm. **E)** Detected glutamate levels in presence of L-glutamate (in green, n = 4 slices) and D-glutamate (in magenta, n = 5 slices). **F)** Quantification of detected iGluSnFR3 fluorescence levels in presence of L-glutamate and D-glutamate. E: data are displayed as mean ± SEM. F: data are displayed as median and the interquartile range (25th to 75th percentiles), whiskers reach to the minimum and maximum values of the distribution; Two-way ANOVA followed by Tukey post-hoc test. Schematic illustrations are created with BioRender.com.

### Cellular *in vivo* imaging of glutamate dynamics using two-photon microscopy

To evaluate whether the GluTrooper line maintains sufficient signal brightness and dynamic range comparable to AAV-mediated expression [3], we employed *in vivo* two-photon microscopy (2PM) to observe glutamate dynamics at cellular resolution in living animals. A cranial window was implanted over the right barrel cortex. After a two-week recovery period, functional imaging was performed in awake, head-fixed mice during air-puff stimulation of the left whisker pad (**Fig. 3A**). As we observed in our *ex vivo* recordings, distinct patches within the neuropil changed their fluorescence levels in spontaneous and evoked conditions (**Fig. 3B-C and Suppl. Video 4**). Spatiotemporal analysis, across detected patches, revealed a robust increase in iGluSnFR3 ΔF/F upon stimulation (**Fig. 3D-E**). Frequency distribution analysis indicated that most iGluSnFR3 %ΔF/F values were distributed around 6% during image acquisition (**Fig. 3F**). Statistical comparison of pooled data from all animals and ROIs confirmed a significant increase in iGluSnFR3 %ΔF/F during stimulation compared to baseline periods (92 detected patches; Baseline vs Peak p<0.0001, **Fig. 3G**).

**Figure 3.**
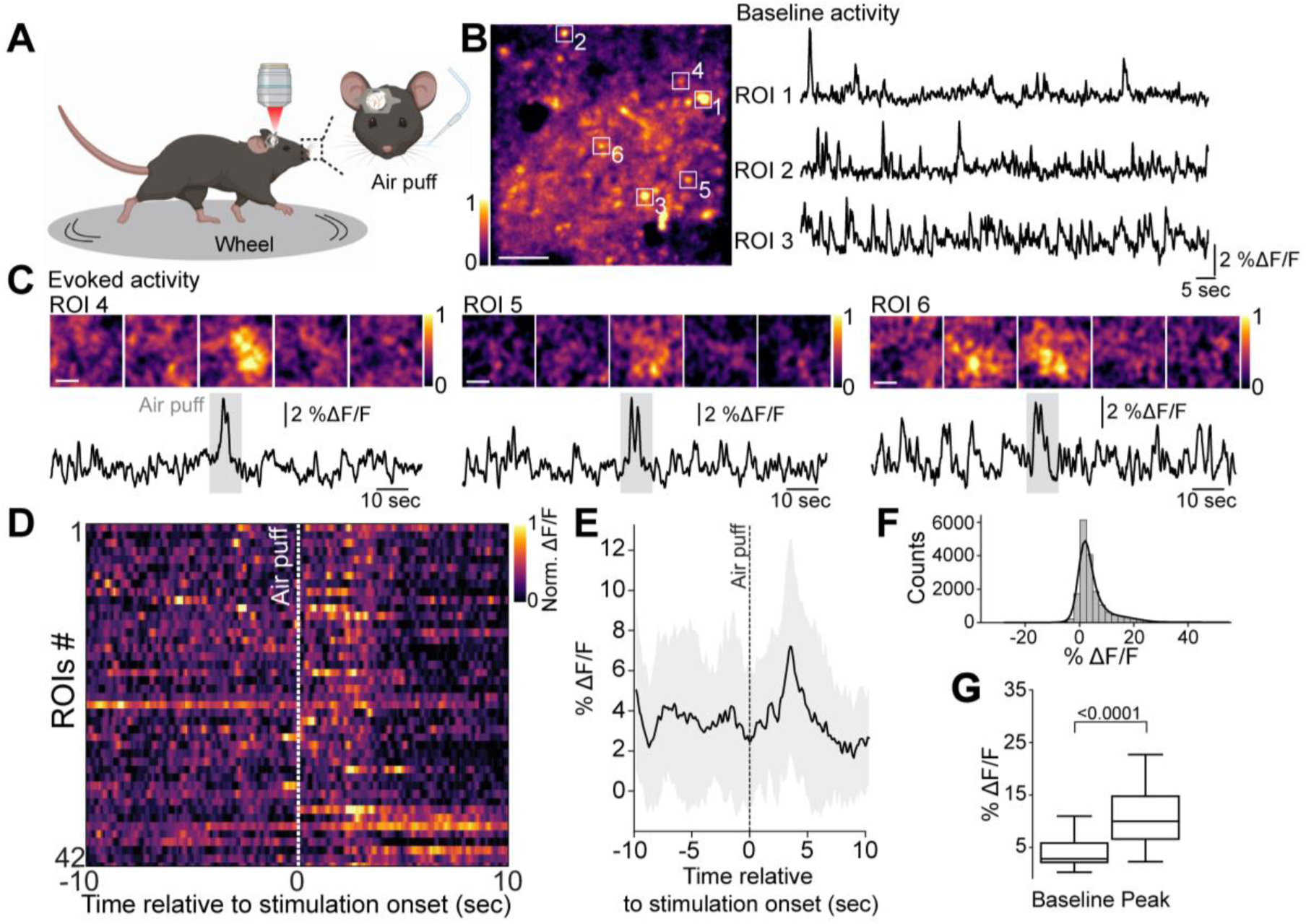
*In vivo* imaging of cellular glutamate dynamics using two-photon microscopy. **A)** Experimental setup schematic and stimulation paradigm. **B)** Representative micrographs and three example ROIs during baseline activity; Scale bar = 10 µm. **C)** Three example ROIs during stimulation period. Note the elevation in fluorescence during stimulation (grey bar); Scale bars = 1µm. **D)** Raster map of analyzed ROIs color-coded by iGluSnFR3 %ΔF/F intensity. Displayed ROIs were detected from the example field of view shown in B. **E–F)** Distribution of iGluSnFR3 response magnitudes during stimulation (**E**) and across the entire experiment. **G)**. Comparison of iGluSnFR3 %ΔF/F between baseline and peak response periods (92 ROIs across 4 mice: p < 0.0001; Wilcoxon matched-pairs signed rank test). E: Data are presented as mean ± SD; G: Data are displayed as median and the interquartile range (25th to 75th percentiles), whiskers reach to the minimum and maximum values of the distribution. Schematic illustrations are created with BioRender.com.

### Wide field cortical glutamate dynamics in awake head-fixed mice

To capture cortex-wide glutamate dynamics *in vivo*, we employed widefield mesoscale imaging (**Fig. 4A**). Since neuronal activity is tightly coupled with hemodynamic responses, which can significantly influence fluorescence-based signals, we used a multispectral illumination approach with a three-wavelength LED system (**Fig. 4A**). This setup allowed simultaneous recording of iGluSnFR3 together with changes in oxyhemoglobin (ΔHbO) and deoxyhemoglobin (ΔHbR) (**Fig. 4B**). Hemodynamic changes can confound fluorescence signals as hemoglobin strongly absorbs light at the excitation and emission wavelengths of GFP-based sensors [16]. These fluctuations in blood oxygenation can alter fluorescence signals independent of actual glutamate dynamics, complicating data interpretation [16, 17]. To address this, we extracted the hemodynamic components and corrected the glutamate signal accordingly. This approach allows for accurate measurement of glutamate dynamics while simultaneously providing a readout of neurovascular coupling, demonstrating that the GluTrooper mouse line is a suitable tool for such investigations. Sensory-evoked glutamate release was elicited using a 10-second tactile stimulus applied to the left whisker pad via a custom Arduino-controlled system (**Fig. 4A and Suppl. Video 5**). Glutamate responses were quantified in the right primary somatosensory cortex (S1), contralateral to the site of stimulation, with the corresponding region in the left hemisphere serving as a endogenous control. Under baseline conditions, spontaneous rate of glutamate events (**Fig. 4C left**) and amplitude (**Fig. 4C right**) were comparable between S1 cortices. However, whisker stimulation produced a significant increase in both the rate of glutamate events (**Fig. 4D left**; p = 0.006) and their amplitude (**Fig. 4D right**; p = 0.04) in the right S1, confirming stimulus specificity of the observed iGluSnFR3 dynamics. To further dissect the relationship between neural and vascular activity, we examined the temporal profile of iGluSnFR3 alongside ΔHbO and ΔHbR during baseline and stimulated conditions on the corrected signals (**Fig. 4E-F**). Whisker stimulation evoked a robust increase in iGluSnFR3 signal (>11% ΔF/F; p = 0.007), alongside with a significant rise in ΔHbO (>12% ΔHbO; p = 0.006) and a reciprocal drop in ΔHbR (<3% ΔHbR; p = 0.006; **Fig. 4E-F**). Notably, temporal alignment of signal peaks demonstrated that glutamate elevations were followed by vascular changes, with iGluSnFR3 reaching its maximum ∼0.56 s after stimulation onset, followed by ΔHbO and ΔHbR peaks at 2.83 s and 2.94 s, respectively (**Fig. 4G**), indicating physiologically relevant temporal pattern of cerebral blood flow regulation. Moreover, correlation analyses during spontaneous activity revealed a strong positive association between glutamate fluctuations and ΔHbO, and an inverse relationship with ΔHbR (**Fig. 4H left**); positive and negative correlations that became stronger upon whisker stimulation (**Fig. 4H right**). These results establish the GluTrooper as the first transgenic model for neurovascular coupling research using a primary neurotransmitter sensor. Although transgenic iGluSnFR expression was previously demonstrated for sensory mapping [18, 19], no prior study has combined stable genetic expression of a glutamate sensor with hemodynamic imaging to investigate neurovascular coupling. By directly measuring glutamate, GluTrooper enables precise temporal correlation between synaptic neurotransmission and vascular responses under both physiological and pathological conditions.

**Figure 4.**
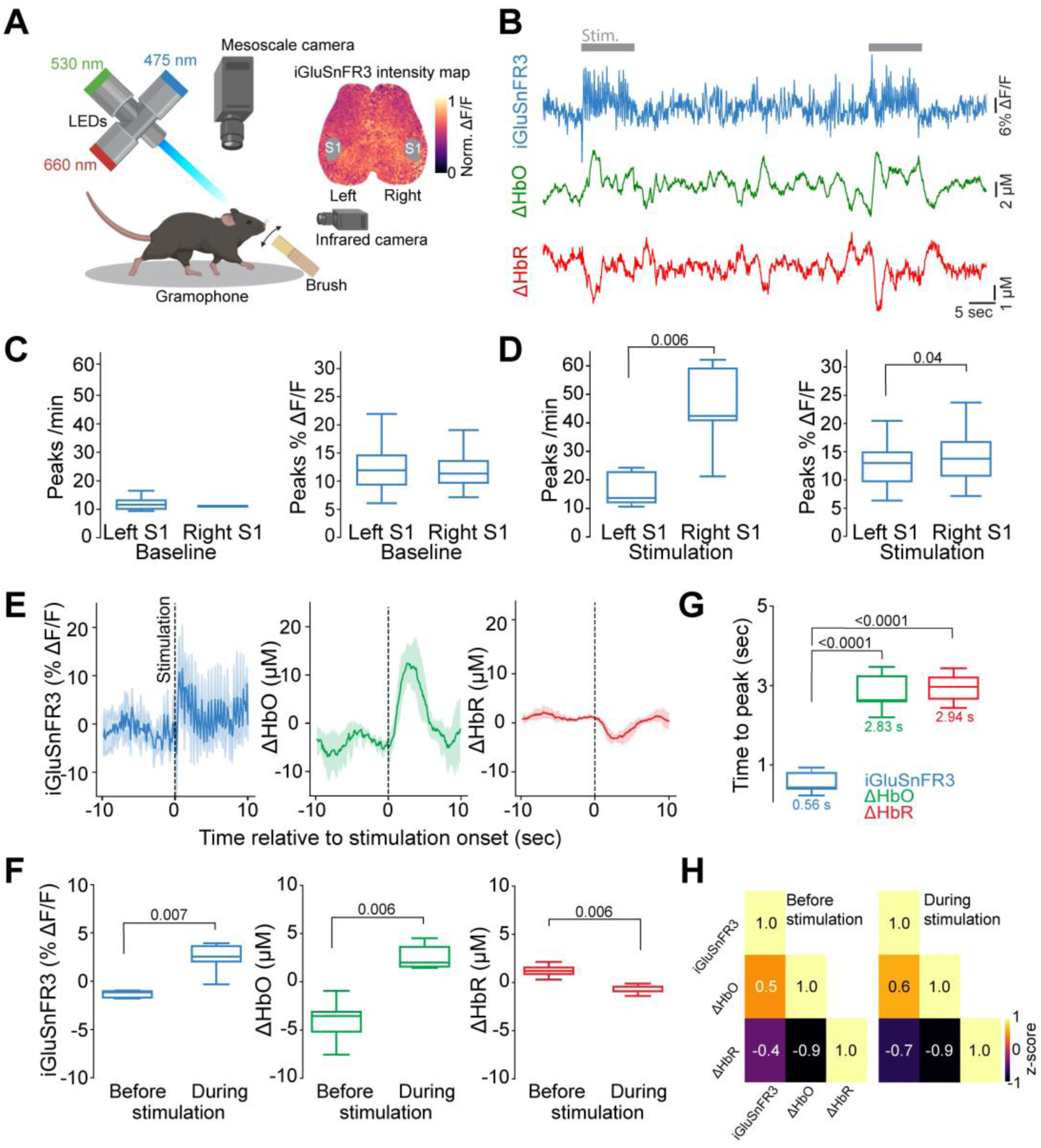
Wide field cortical glutamate dynamics in awake head-fixed mice. **A)** Schematic of the widefield imaging setup and experimental paradigm (on the left), with a representative widefield fluorescence image highlighting the bilateral somatosensory cortex (S1) regions of interest (on the right). **B)** Representative iGluSnFR3 (in blue), ΔHbO (in green) and ΔHbR (in red) traces detected in spontaneous and evoked conditions (stimulation periods are represented by the grey lines). **C)** Number of peaks rate (peaks/min, on the left) and amplitude of detected peaks (%ΔF/F, on the right) during baseline conditions. **D)** Number of peaks rate (peaks/min, on the left; p: 0.006, Student’s t-test) and amplitude of detected peaks (%ΔF/F, on the right; p = 0.04, Mann Whitney test) during whisker stimulation. **E)** Simultaneous recording of iGluSnFR3 and hemodynamic responses, along with stimulus timeline following stimulation onset. **F)** iGluSnFR3 signaling intensity and hemodynamic parameters during pre-stimulus baseline vs. active stimulation periods (iGluSnFR3 %ΔF/F: p = 0.007; ΔHbO: p = 0.006; ΔHbR: p = 0.006, Student’s t-test). **G)** Time to peak of iGluSnFR3 %ΔF/F, ΔHbO and ΔHbR from stimulation onset (iGluSnFR3 vs ΔHbO: p = <0.0001; iGluSnFR3 vs ΔHbR: p = <0.0001, n animals = 5, One-way ANOVA followed by Tukey’s post-hoc test). **H)** Correlation matrix of iGluSnFR3, ΔHbO, and ΔHbR during baseline and stimulation, showing increased coupling between glutamate and hemodynamic signals during stimulation (see color scale for z-score values). C-D-G-F: Data are displayed as median and the interquartile range (25th to 75th percentiles), whiskers reach to the minimum and maximum values of the distribution. E: Data are presented as mean ± 95% confidence interval. Schematic illustrations are created with BioRender.com.

### *In vivo* bilateral multimodal imaging of neurovascular coupling during cortical spreading depolarizations

After assessing glutamate dynamics in spontaneous and evoked conditions, we utilized the GluTrooper line for the observation of cortical spreading depolarizations (CSDs) [20], a pathological wave of near-complete neuronal and glial depolarization that propagates slowly across the cerebral cortex. This phenomenon is characterized by widespread glutamate release and a breakdown of ion homeostasis, leading to prolonged neuronal silencing and often resulting in impaired vascular responses in injured brain tissue [21]. To trigger a CSD in the left hemisphere a small burr hole was drilled over the left occipital cortex and a 1M KCl^-^ solution was topically applied to the exposed cortical surface (**Fig. 5A**). Spatiotemporal glutamate dynamics during CSDs were quantified over multiple regions of interests (ROIs) distributed along the caudal-rostral axis in both hemispheres (**Fig. 5B**). Upon KCl^-^ application, CSD was elicited and propagated across the left hemisphere, causing a more then 10-fold increase in iGluSnFR3 %ΔF/F across all ROIs indicative of massive synaptic glutamate release (**Fig. 5C-D and Suppl. Video 6**; Baseline vs CSD). Spatiotemporal mapping of the depolarization wave yielded a propagation velocity of 3.63 mm/min (**Fig. 5E)** and a significantly greater area under the curve in the ipsilateral hemisphere compared to the contralateral side (**Fig. 5F**; p = 0.0009). By contrast, examination of ROIs in the contralateral hemisphere revealed modest glutamate fluctuations during CSD propagation (**Fig. 5G)** with the strongest increase observed in the most frontal ROI (∼50% of iGluSnFR3 % ΔF/F; **Fig. 5H**), suggesting an indirect contralateral response to the unilateral perturbation. To assess concurrent cerebral hemodynamic changes, we simultaneously measured ΔHbO and ΔHbR in the same bilateral ROIs across three distinct temporal windows: Before CSD, CSD onset, and post-CSD [22, 23] (**Fig. 5I**). In the ipsilateral hemisphere, ΔHbO significantly increased post-CSD (**Fig. 5J**; p = 0.001), consistent with the transient hyperemic response and perfusion-driven oxygen delivery described in previous studies [24], while ΔHbR is significantly reduced in post-CSD phase (**Fig. 5K**, p = 0.004). In contrast, the contralateral hemisphere exhibited no significant changes in ΔHbO or ΔHbR, suggesting that absolute hemodynamic alterations were largely confined to the affected hemisphere (**Fig. 5L-N**). Together, these findings underscore the utility of the GluTrooper reporter line for high-resolution, multimodal imaging of excitatory neurotransmission and vascular dynamics during CSD. Most importantly, this approach enables a more direct, non-invasive assessment of neurovascular coupling under pathological conditions, offering a viable alternative to traditional invasive techniques.

**Figure 5.**
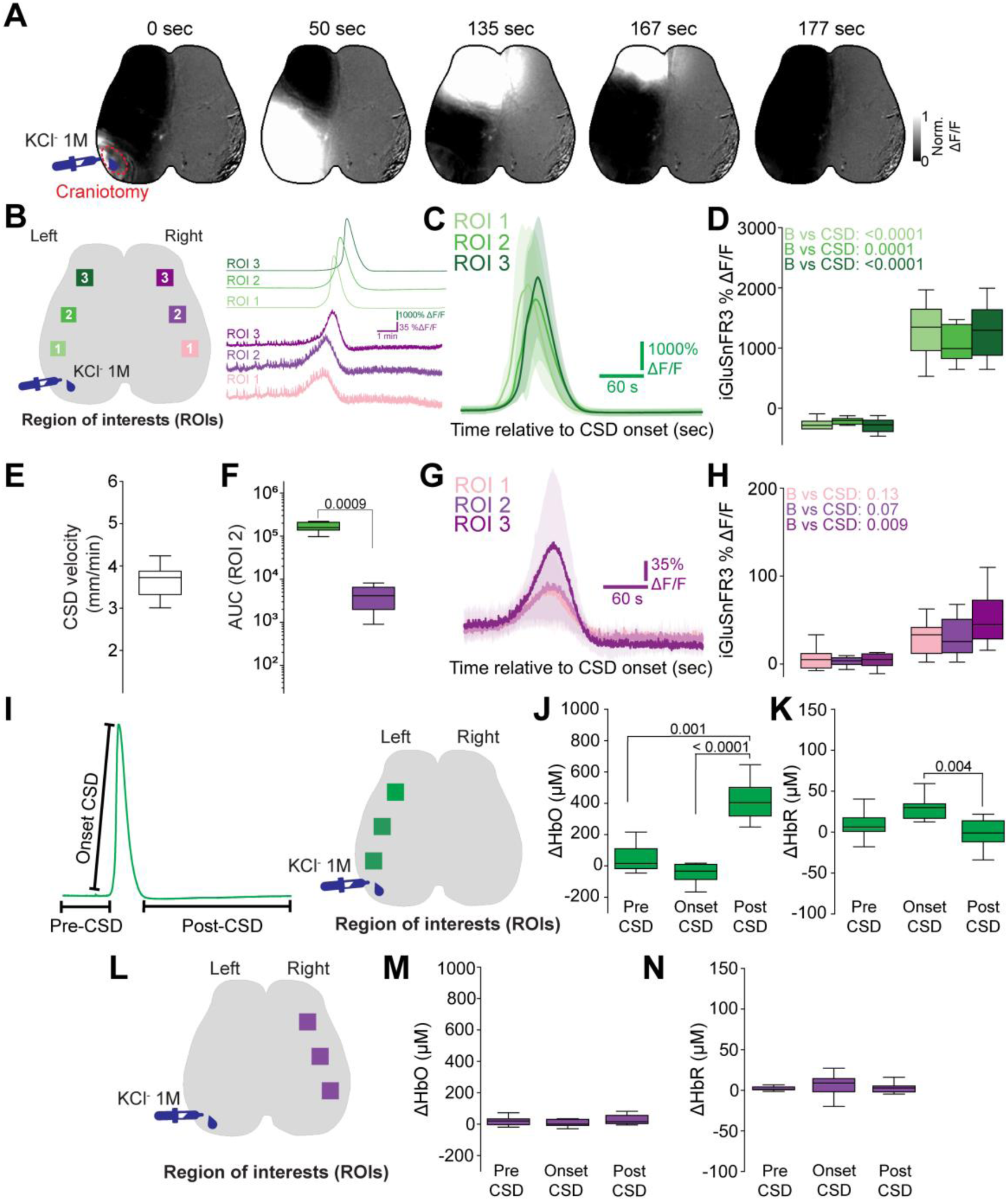
*In vivo* bilateral multimodal imaging of neurovascular coupling during cortical spreading depolarizations. **A)** Representative image series showing KCl^-^ application site and the progression of the CSD in the left hemisphere. **B)** Schematic of bilateral regions of interest (ROIs) and corresponding extracted iGluSnFR3 dynamics **C)** Glutamate dynamics (%ΔF/F iGluSnFR3) in the left ROI aligned to CSD onset. **D)** Comparison of mean iGluSnFR3 %ΔF/F in the left ROIs during baseline (B) and CSD (ROI1 and 3: p <0.0001; ROI2: p = 0.0001, Two-way ANOVA followed by Tukey’s post hoc test). **E)** Quantification of CSD propagation velocity. **F)** Area under the curve (AUC) of glutamate transients in bilateral ROI2 (left vs right: p = 0.0009, Student’s t-test). **G)** Glutamate dynamics (%ΔF/F iGluSnFR3) in the right ROI aligned to CSD onset. **H)** Comparison of mean iGluSnFR3 %ΔF/F in the right ROIs during baseline (B) and CSD (ROI3, p: 0.009, Two-way ANOVA followed by Tukey’s post hoc test). **I)** Schematic representing the analyzed CSD phases (on the left) and distribution of ROIs in the left hemisphere. **J-K)** Quantification of ΔHbO (J) and ΔHbR (K) across CSD phases in the left hemisphere (For ΔHbO, Pre-CSD vs Post-CSD: p = 0.001; Onset-CSD vs Post-CSD: p < 0.0001; For ΔHbR, Onset-CSD vs Post-CSD: p = 0.004; Friedman test followed by Dunn’s multiple-comparisons test). **L)** Schematic of the ROIs distributed on the right hemisphere. **M-N**) Quantification of ΔHbO (J) and ΔHbR (K) across CSD phases in the right hemisphere. C-G: Data are presented as mean ± 95% confidence interval; D-H-J-K-M-N: Data are displayed as median and interquartile range (25th–75th percentiles), whiskers reach to the minimum and maximum values of the distribution, n = 6 mice.

### Widefield mapping of contralateral disinhibition following CSDs

After observing widespread glutamate fluctuations during CSDs, we next asked whether the GluTrooper line is sensitive enough to detect subtle alterations in cortical neuronal network dynamics, such as contralateral disinhibition, a phenomenon linked to impaired interhemispheric inhibition following stroke and brain injury [25]. To investigate this, we selected one ROI in the center of each hemisphere and analyzed iGluSnFR3 fluorescence traces from the widefield recordings, allowing continuous monitoring of glutamate transients before and after KCl^-^-induced CSD (**Fig. 6A**). Analysis of the left hemisphere revealed alterations in glutamate dynamics following CSD (**Fig. 6B**). Although the amplitude of detected iGluSnFR3 events did not change significantly (**Fig. 6C left**; p = 0.07), a significant reduction in peak rate was observed post-CSD (**Fig. 6C right**; p = 0.01). These results are in line with the extensive literature focused on CSD, representing the strong long-lasting cellular depression that follows this phenomenon [26–29]. In contrast, the contralateral hemisphere exhibited a markedly different response profile (**Fig. 6D**). Post-CSD, event amplitudes remained stable (**Fig. 6E left**; p = 0.38), however, we observed a significant increase in the rate of glutamate events following CSD (**Fig. 6E right**; p = 0.01), indicative of contralateral disinhibition [25, 30]. These findings demonstrate that the GluTrooper enables non-invasive optical detection of rapid, transient glutamate events and interhemispheric compensatory activity following CSD. While previous wide-field imaging studies, using Thy1-GCaMP6s mice, successfully mapped large-scale network connectivity changes after cortical injury [31], calcium indicators provide only indirect measures of neuronal activity and cannot resolve neurotransmitter-specific dynamics. By directly reporting extracellular glutamate with high temporal precision, GluTrooper reveals glutamatergic mechanisms underlying network reorganization that remain inaccessible to calcium-based reporters

**Figure 6.**
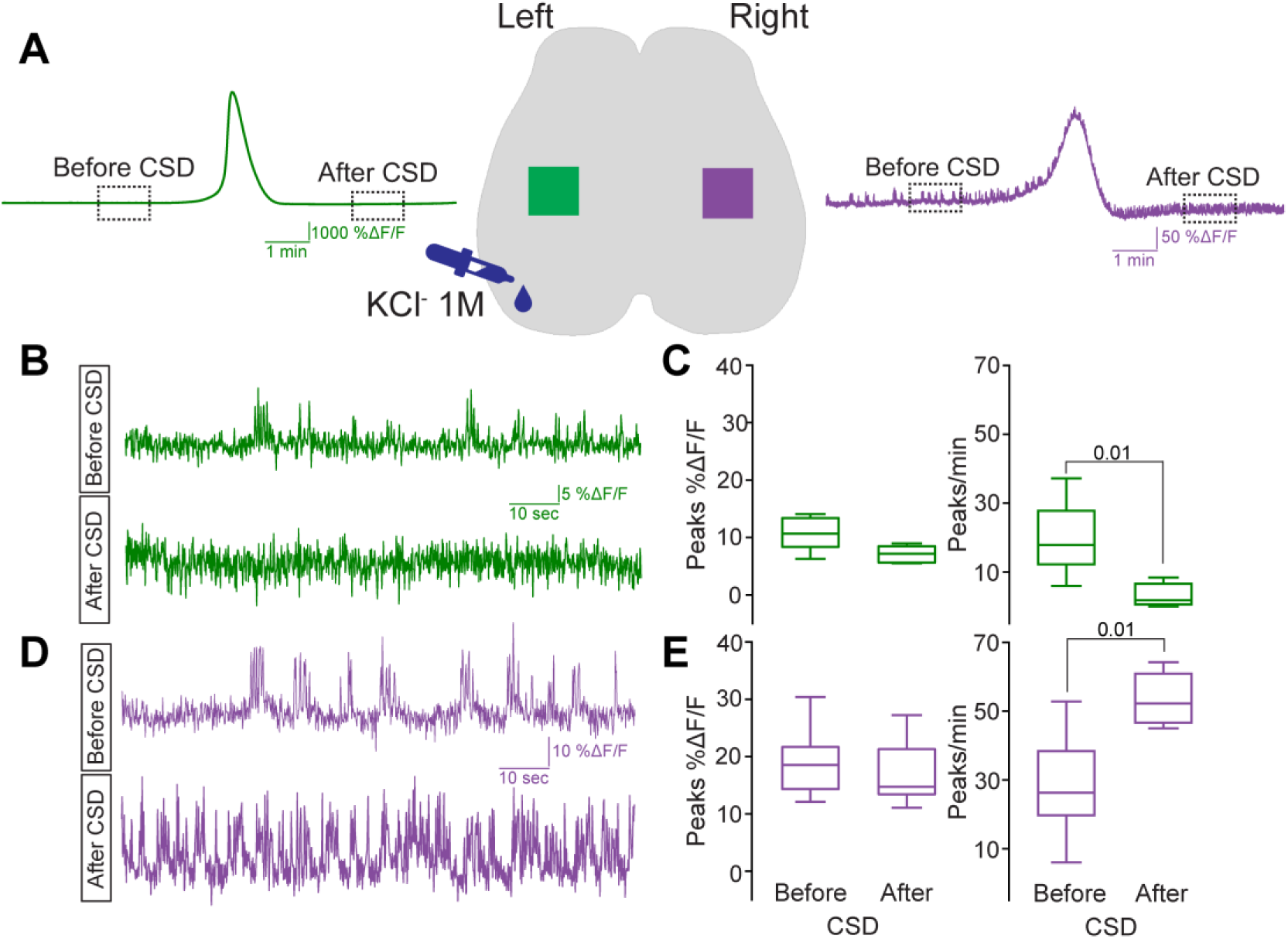
Widefield mapping of contralateral disinhibition following CSD. **A)** Schematic representing the selected bilateral ROIs with representative iGluSNFR3 fluctuations detected from the left ROI (in green on the left) and right ROI (in purple on the right). **B)** Inserts of the iGluSnFR3 traces detected from the left ROI before and after CSD. **C)** Quantification of peaks amplitude (on the left; %ΔF/F p = 0.07) and peaks rate (on the right, peaks/min; p = 0.01) detected from the left ROI before and after CSD. **D)** Inserts of the iGluSnFR3 traces detected from the right ROI before and after CSD. **E)** Quantification of peaks amplitude (on the left; %ΔF/F p = 0.38) and peaks rate (on the right, peaks/min; p = 0.01) detected from the right ROI before and after CSD. C-E: Data are displayed as median and interquartile range (25th–75th percentiles), whiskers reach to the minimum and maximum values of the distribution; n = 6 mice. C-E Student’s t-test.

## Discussion

Here, we described the development and characterization of the GluTrooper, a novel ROSA26 knock-in mouse model for Cre-dependent expression of the glutamate sensor iGluSnFR3 [3–6]. We demonstrate that this transgenic approach overcomes key limitations of viral delivery methods and enables stable, long-term monitoring of glutamatergic signaling across diverse experimental paradigms. Using widefield imaging in awake mice, we show that GluTrooper provides high-fidelity detection of both spontaneous and evoked glutamate dynamics, directly measuring the primary excitatory neurotransmitter and thereby validating glutamate’s role in neurovascular coupling and interhemispheric communication during cortical spreading depolarization, mechanisms previously inferred from calcium-based reporters [23, 31, 32].

### The GluTrooper addresses technical limitations of current glutamate imaging approaches

Traditional methods of glutamate imaging have largely relied on viral vector-based delivery of sensors such as iGluSnFR [3, 4], which, although effective, often result in inconsistent expression levels, limited spatial spread, and cytotoxic effects due to high overexpression in a subset of neurons [7, 33]. Moreover, the transfection requires a surgically invasive procedure, which carries potential risks for the animal and may alter the cortical microenvironment. By leveraging the ROSA26 locus and the strong, ubiquitous CAG promoter, the GluTrooper overcomes these limitations, offering a robust, reproducible, non-toxic and non-invasive alternative for brain-wide glutamate imaging. Crossing our novel reporter line with Emx1-Cre mice, we observed efficient recombination and expression across the cortex, as well as in the hippocampus and olfactory bulb [12, 13]. The stability of sensor expression over time further enhances the utility of this model for chronic imaging paradigms, including longitudinal monitoring of synaptic function, learning, and plasticity. To verify its functionality, we first conducted *ex vivo* glutamate imaging on acute brain slices, demonstrating that iGluSnFR3 expressed in our novel reporter line retains the same glutamate selectivity as the virally delivered sensor [15], enabling the reliable detection of both spontaneous and evoked glutamate dynamics.

### The GluTrooper enables multimodal imaging of interhemispheric glutamate dynamics

To further validate the functionality of our novel glutamate reporter line, we employed widefield glutamate imaging in awake, head-fixed mice. We confirmed that iGluSnFR3 expression was sufficiently bright and stable to permit high signal-to-noise recordings of both spontaneous and evoked glutamatergic dynamics. In the barrel cortex, sensory stimulation reliably evoked glutamate transients with high temporal precision and regional specificity. These recordings also revealed a tight coupling between cortical activation and glutamate release, reinforcing the validity of this sensor for monitoring specific neuronal output rather than general network excitability. During our experimental approaches, we observed that glutamate transients temporally preceded hemodynamic responses, including changes in ΔHbO and ΔHbR, revealing the precise timing between sensory-evoked neurotransmitter release and vascular responses. By measuring glutamate as the primary excitatory neurotransmitter rather than downstream calcium signals, iGluSnFR3 enables improved temporal resolution for characterizing neurovascular coupling. It also opens up new doors for investigating the role of excitatory signaling in regulating vascular responses under physiological and pathophysiological conditions. Beyond physiological validation, we tested our novel reporter line in the context of cortical spreading depolarization, a hallmark of several neurological conditions including stroke, traumatic brain injury, and migraine [21]. Using three different LEDs, we were able to simultaneously track glutamate dynamics and hemodynamic responses across the cortex of both hemispheres. We selected three distinct phases, baseline, CSD onset, and post-CSD [22], and extracted glutamate and vascular dynamics from homotypic regions of interest in both hemispheres. As expected, CSD evoked a pronounced increase in glutamate concentration in the ipsilateral cortex, coupled with a complex hemodynamic response characterized by an initial hypoperfusion followed by delayed hyperemia. Moreover, following the observation of spontaneous glutamate dynamics before and post- CSD, we quantified a significant increase in the frequency of spontaneous contralateral glutamate events during the post-CSD phase. This data shows the sensitivity of the GluTrooper in observing contralateral disinhibition, phenomenon that can be triggered by unilateral lesion or brain injury [34, 35]. The ability to observe this phenomenon non-invasively, through direct imaging of glutamate release, provides a unique window into interhemispheric network dynamics during cortical perturbations. It also raises important questions about the role of contralateral plasticity in recovery or maladaptation following focal insults. Taken together, our findings establish GluTrooper, our novel glutamate reporter line as a reliable, flexible, and scalable tool for investigating glutamatergic signaling in acute brain slices and in awake, walking animals. This model enables chronic, high-fidelity imaging of glutamate transients across a wide range of spatial and temporal scales, overcoming many of the technical limitations inherent to viral delivery methods. By allowing for longitudinal monitoring of cortical function, sensory processing, and interhemispheric communication, GluTrooper opens new opportunities for studying circuit-level excitatory dynamics under both physiological and pathophysiological conditions.

### Limitations of the study

While our novel reporter line provides a robust and versatile platform for the observation of glutamate dynamics, several limitations should be considered. First, although the Cre-dependent design allows for targeted expression of iGluSnFR3, the time required to breed animals is longer than what is required to obtain viral expression following intracortical injection. Second, the uniform and widespread expression driven by the CAG promoter may dilute spatial contrast in densely interconnected networks, complicating interpretation in areas where fine spatial resolution of glutamate dynamics is critical. Another consideration is the biological impact of long-term iGluSnFR expression, particularly under conditions of chronic imaging or repeated stimulation. Although we did not observe cytotoxicity or altered neuronal responses in widefield or two-photon imaging, subtle alterations in synaptic physiology or glutamate uptake cannot be fully excluded. Previous studies have suggested that glutamate sensors can alter glutamate clearance or buffering, especially at high expression levels [36]. The knock-in design of the GluTrooper mitigates this risk by driving moderate, consistent expression levels, yet future studies should directly assess synaptic function and plasticity in labeled versus unlabeled neurons to rule out undesired aberrant processes.

In summary, the GluTrooper mouse line is a significant methodological advance for investigating neuronal dynamics *in vivo*. Most importantly, it allows the monitoring of the first messenger, glutamate, whereas current approaches primarily monitor the second messenger, i.e. the rise in intracellular neuronal calcium. Furthermore, compared to current viral-based approaches to imaging glutamate in the brain, the GluTrooper mouse line offers long-term sensor stability, extensive spatial coverage, and genetic specificity. This makes the GluTrooper mouse line a unique tool for the longitudinal observation of glutamate dynamics, minimizing technical artefacts that could compromise data quality. Using this model, researchers can monitor glutamatergic neurotransmission in the whole brain with exceptional reproducibility without needing to use invasive delivery methods or risking the aberrant processes associated with AAV-based strategies. Furthermore, its compatibility with various Cre-driver lines establishes the GluTrooper mouse line as a valuable resource for future studies focusing on specific cell types or brain regions. This will further expand our understanding of glutamatergic neurotransmission in healthy brains and pathological contexts such as excitotoxic injury and neurodegeneration.

### Material and methods Animals

RCL-iGluSnFR3 x Emx1-Cre mice were used for all experiments. Animals were group-housed under pathogen-free conditions and bred in the animal facility of the Center for Stroke and Dementia Research. Food and water were provided ad libitum under controlled environmental conditions (21 ± 1 °C, 12/12-h light/dark cycle). All experimental procedures complied with the ARRIVE guidelines, the National Guidelines for Animal Protection (Germany), and were approved by the regional Animal Care Committee of the Government of Upper Bavaria.

### Immunohistochemistry

Mice were deeply anesthetized and transcardially perfused with ice-cold 4% paraformaldehyde (PFA) in 0.1 M phosphate buffer. Brains were post-fixed in the same fixative for 24 hours at 4°C, then 100- μm sagittal sections were collected with a Leica vibratome (Leica Biosystems, Germany). Sections were incubated with primary antibody (goat anti-GFP; Thermo Fisher Scientific, #600-101-215M) diluted in blocking solution overnight at 4°C to observe iGluSnFR3-expressing cells. After three PBS-T (0.05% Tween 20) washes, tissue was incubated with species-matched Alexa Fluor 488-conjugated secondary antibody (donkey anti-goat IgG (H+L); Thermo Fisher Scientific, #A-11055) diluted 1:500 in PBS-T and DAPI (Invitrogen, #D1306; 1:1000). Then, slices were washed 3 times for 20 minutes each prior to mounting.

### Confocal microscopy

Confocal imaging was performed using a Leica Stellaris 5 system (Leica Microsystems, Wetzlar, Germany) equipped with 40× (NA 1.30) and 63× (NA 1.40) oil-immersion objectives. Z-stacks were acquired at 2,048 × 2,048-pixel resolution with 0.65 µm z-intervals. Four neuroanatomically distinct regions (somatosensory cortex, hippocampus, olfactory bulb, and cerebellum) were systematically imaged in both 3- and 12-month-old cohorts to evaluate age-dependent expression profiles. Post-acquisition processing utilized Leica LAS X software (v3.7.4, Leica Microsystems) to generate maximum intensity projections from raw z-stacks. For cross-regional comparisons, cerebellar iGluSnFR3 signals served as an internal control due to its lack of transcriptional factor Emx1. Comparison of fluorescence intensity between the different regions and age was done using Python (v3.9) with Scikit-package.

### Chronic widefield window implantation

Multimodal widefield imaging was performed to simultaneously monitor extracellular glutamate dynamics and intrinsic optical signals (IOS) reflecting cortical hemodynamics, using a custom-built mesoscale imaging system. Cranial window preparation was conducted three days prior to imaging sessions. Animals were anesthetized with isoflurane (5% for induction, 2% for maintenance) in a gas mixture of 70% nitrous oxide and 30% oxygen, and positioned in a prone orientation in a stereotaxic frame (Stoelting, Europe, #51501). After removal of scalp and connective tissue, the skull was coated with a thin layer of transparent dental cement (Quick Base S398, L-Powder clear S399, Universal Catalyst S371, Parkell C&B metabond, USA). A curved crystal glass coverslip (Crystal Skull, LabMaker, USA) was then affixed over the intact skull using the same cement, and a custom head-fixation frame was attached around the window. Animals recovered for a minimum of 48 hours before imaging.

### Chronic cranial window implantation

Mice were anesthetized via intraperitoneal injection of a mixture containing medetomidine (0.5 mg/kg), midazolam (5 mg/kg), and fentanyl (0.05 mg/kg; MMF). A craniotomy was made over the right barrel cortex (from bregma −1.1 mm, lateral +3.3 mm). The bone flap and dura were carefully removed, and a circular 3 mm glass coverslip was placed over the exposed cortex and sealed to the skull with Vetbond^®^. To enable stable head fixation during imaging, a titanium head plate (Femtonics Ltd., Budapest, Hungary) was affixed to the skull using UV-curing dental cement. After surgery, anesthesia was reversed with atipamezole (0.5 mg/ml), naloxone (3 mg/ml), and flumazenil (5 mg/ml; ANF) administered intraperitoneally. Mice were kept in a 32°C heating chamber overnight to ensure complete recovery and were given two weeks to recover before *in vivo* imaging.

### Awake widefield mesoscale imaging and sensory stimulation

Mice were briefly anesthetized with 4% isoflurane to allow head fixation onto a Gramophone-style holder (Grammophone-1, Femtonics, Hungary) using the custom head frame. Sensory stimulation was delivered to the left whisker pad using a custom-built, Arduino-controlled whisker stimulator oscillating vertically at 2 Hz. Each session included a 4-minute habituation period followed by a 4-minute imaging protocol consisting of a 50-second baseline and four stimulation blocks (10 s stimulation, 20 s rest), concluding with a final rest period. Infrared video was recorded concurrently using a Basler camera. To prevent visual interference from the imaging system, a 3D-printed eye shield was placed over the animal’s eyes. Following video correction and ΔF/F extraction from the selected regions of interest, signals were analyzed using SciPy (Python). The left and right somatosensory cortices were identified using the Allen Brain Atlas (Allen Institute, USA) as reference. After baseline correction, glutamate events were detected by calculating the mean trace and applying a threshold of 1.5 standard deviations to ensure that only true fluctuations were considered as events. Event detection was performed using SciPy (Python), and statistical differences in detected peaks during baseline and stimulation in both somatosensory cortices were compared using GraphPad Prism. Bulk changes in iGluSnFR3, ΔHbO, and ΔHbR were assessed by comparing the 10 seconds before and after stimulation onset. The time to peak of these bulk changes was measured from the initiation of stimulation to the peak of the signals.

### Cortical Spreading Depolarization

To observe CSD, mice were administered medetomidine (0.05 mg/kg, i.p.) five minutes prior to induction with 5% isoflurane delivered in a gas mixture of 70% nitrous oxide and 30% oxygen. After 1 minute, animals were placed in a stereotaxic frame, and isoflurane was gradually reduced to 1.5% for 140 seconds, followed by 0.75% for an additional 2 minutes to achieve a stable anesthetic state suitable for imaging. A small craniotomy was performed approximately one hour prior to imaging, positioned caudo-laterally to the primary imaging window, and filled with sterile phosphate-buffered saline (PBS). Following a 2-minute baseline period, CSD was induced by replacing the PBS with 1M KCl for 10 seconds, after which PBS was promptly reintroduced. Imaging continued for an additional 10 minutes, resulting in a total recording time of 12 minutes. Successful induction of CSD was verified through IOS monitoring. Immediately after imaging, animals were sacrificed. To quantify changes in glutamate dynamics, the mean ΔiGluSnFR3, ΔHbO, and ΔHbR signal from all defined regions of interest was calculated and compared between the baseline and CSD periods using GraphPad Prism.

### Pre-processing of widefield imaging data analysis

Data acquisition and preprocessing were conducted using methodologies partially described previously [31, 37], supplemented by newly developed analytical approaches. All image processing and analyses were performed using MATLAB (Mathworks, R2016b), with additional custom scripts written in Python (version 3.11.3) for extended analyses and visualization tasks. Briefly, acquired images underwent motion correction and alignment, background fluorescence subtraction, and normalization of fluorescence signals to the mean intensity across each recording for individual pixels (ΔF/F).

### *In vivo* Two-Photon microscopy

Two-photon glutamate imaging was performed two weeks following cranial window implantation to allow for complete tissue recovery. Experiments were conducted using an Atlas two-photon microscope (Femtonics Ltd., Budapest, Hungary) equipped with a Chameleon Ultra tunable laser (Coherent Inc, USA) and a 16× water immersion objective (Nikon, NA 0.8, Japan). The iGluSnFR3 sensor was excited at 940 nm with fluorescence emission collected between 500-550 nm. Average laser power administered was ≤ 40 mW to prevent tissue damage. Awake mice were positioned on a circular head-fixation platform (Gramophone, Femtonics Ltd, Hungary) and allowed to habituate for 10 minutes prior to recording. Time-lapse imaging was acquired at 9.8 Hz (1000 × 1000 pixels, 50µm x 50µm) at cortical depths between 150-200 μm below the surface. Each recording session began with a 1-minute baseline period, followed by manually controlled whisker stimulation delivered via Picospritzer (Parker-Hannifin Corporation) at ⁓5 Hz frequency with 80 ms pulse duration for 10 seconds. All image processing and quantitative analyses were conducted after image acquisition.

### Two-photon glutamate imaging analysis

Motion correction was first performed using the NoRMCorre [38] to reduce movement-related artifacts in the imaging data. After stabilization, time-lapse sequences were imported into Fiji, where regions of interest corresponding to glutamate-release areas were manually identified and fluorescence traces were extracted from each patch. For each labeled region, changes in ΔF/F glutamate levels were quantified by comparing the five seconds preceding stimulation with the peak detected during whisker stimulation. All statistical analyses were conducted using GraphPad Prism to assess differences in glutamate dynamics between experimental conditions.

## Author Contributions

BW and WW generated the mouse. GMC, DPV, SF and NP contributed to the study design. GMC, DPV, PF, SF, FG, JJS, AC and SG contributed to the acquisition of the data with the support of NP, WW, JH, LB. GMC and DPV analyzed the data and performed statistical analysis. GMC, DPV and NP drafted the manuscript.

## Funding

This work was funded by the Solorz-Zak Foundation and the Deutsche Forschungsgemeinschaft (DFG, German Research Foundation) under Germany’s Excellence Strategy within the framework of the Munich Cluster for Systems Neurology (EXC 2145 SyNergy – ID 390857198; NP) and under a project grant to NP (PL 249/24-1; ID 551429233).

The work was also supported by the DFG under the Heisenberg Programme (Project No. 447395247) (to L.F.B.) and under Germany’s Excellence Strategy within the framework of the Munich Cluster of Systems Neurology (EXC 2145 SyNergy—ID 390857198) (to L.F.B.).

## Competing Interests

The authors declare no competing interests.

## Data availability

The data generated during the study are available from the corresponding author upon request.

## Notes

### Competing Interest Statement

The authors have declared no competing interest.

